# NeuroMASS v4: Geometry-Aware Neural Population Dynamics via Learnable Hyperbolic Latent Representations and Biologically-Calibrated Diagonal State Space Models

**DOI:** 10.64898/2026.11.13.766497

**Authors:** Ranveer Singh, Jaideep Varma, Aryan Somanna

## Abstract

The geometry of neural population activity is a fundamental challenge in computational neuroscience. We present NeuroMASS v4, a novel architecture that introduces learnable hyperbolic latent geometry to spike-train co-smoothing on the Neural Latents Benchmark (NLB’21). Our central scientific claim is that the curvature *c* of a Poincaré-ball—learned end-to-end from extracellular spike recordings— constitutes a biologically interpretable geometric signature: significantly positive for ring-manifold motor circuits (MC_Maze, MC_RTT) and near-zero for topologically unstructured frontal circuits (DMFC_RSG). The architecture unifies five principled components: (i) discrete spike-count embeddings with learned neuron-identity vectors, (ii) permutation-equivariant attention-weighted neuron pooling, (iii) a diagonal S4D state space model initialised with a log-uniform frequency prior spanning 2–200 Hz, (iv) a Poincaré-ball projection with learnable scalar curvature, and (v) a Frobenius-bounded session adapter. Training uses masked self-prediction of spike counts with an auxiliary behaviour decoding objective; all experiments use exclusively real DANDI/NLB recordings under a strict fail-stop data policy. We report results across four NLB datasets spanning motor, somatosensory, and cognitive cortices in rhesus macaque, and demonstrate that learned curvature statistically predicts circuit topology.

## 1 Introduction

The neural population code—the collective activity pattern of hundreds of simultaneously recorded neurons—is now accessible to routine measurement via Utah arrays, Neuropixels probes, and ECoG grids. This experimental capability has outpaced our theoretical understanding of how populations encode behavioral variables and how their representational geometry changes across sessions, subjects, and brain regions.

A central observation in motor neuroscience is that M1 and PMd population trajectories during arm reaching lie on approximately ring-shaped manifolds whose angular coordinate encodes reach direction [1, 2]. Such ring manifolds possess intrinsic positive curvature—they cannot be globally isometrically embedded in any Euclidean space without distortion. Every existing neural latent variable model— GPFA [3], LFADS [4], AutoLFADS [5], NDT [6], PopT [7]—uses a Euclidean latent space, necessarily incurring this distortion, limiting interpretability and obscuring the geometric signal encoding circuit organisation.

Hyperbolic geometry natively represents ring and hierarchical structures with low distortion [8, 9]. A Poincaré ball of curvature *c* embeds a discretised ring of *n* elements with distortion *O*(1/*n*) versus *O*(1) in Euclidean space. We make *c* a learnable parameter trained end-to-end from spike data. When *c* converges above zero, the model has discovered hyperbolic structure from data; when *c* → 0, it gracefully recovers Euclidean geometry.

### 1.1 Contributions

- **Learnable Hyperbolic Geometry**. First neural population dynamics model with a learnable Poincaré-ball curvature *c*, correctly derived from the Möbius exponential map at the origin [9]. Round-trip error < 10^−15^.
- **NeuralS4D Timescale Prior**. Log-uniform frequency prior over [2, 200] Hz replacing generic HiPPO initialisation, spanning the theta/beta/gamma oscillatory hierarchy.
- **Permutation-Equivariant Neuron Pooling**. Attention-weighted pooling over learned neuron-identity embeddings; invariant to electrode channel reordering between sessions.
- **Bounded Session Adapter**. Low-rank adapter with Frobenius-norm constraint guaranteeing representational drift ≤ *δ* (*∥z∥*+ 1) for any context.
- **Fail-Stop Real-Data Pipeline**. Exclusively real DANDI/NLB recordings throughout; download failures terminate with diagnostics rather than substituting synthetic data.

## 2 Related Work

### 2.1 Neural Latent Variable Models

GPFA [3] introduced Gaussian-process factor analysis under a linear-Gaussian emission model. LFADS addressed nonlinearity with a bidirectional GRU encoder and sequential GRU generator. AutoLFADS [4] scaled LFADS with population-based Bayesian hyperparameter optimisation. NDT [6] adapted BERT’s masked self-prediction to spike trains. PopT [7] extended this with population-level Transformer cross-attention. All methods embed activity in Euclidean latent spaces, geometrically suboptimal for ring-manifold circuits.

### 2.2 State Space Models for Sequences

The structured SSM family [10] demonstrated HiPPO-initialised recurrences for long-range dependencies with *O*(*T* log *T*) training complexity via FFT convolution. S4D [11] simplified to a diagonal complex *A* matrix. Mamba [12] introduced input-dependent state selection. Our NeuralS4D adapts S4D’s diagonal kernel with a neuroscience-motivated frequency prior, stacking *L* = 4 layers with SiLU gating.

### 2.3 Hyperbolic Geometry in Deep Learning

Poincaré embeddings [8] demonstrated hyperbolic space for hierarchical relational data. Hyperbolic Neural Networks [9] extended this to deep architectures via Möbius transformations. Products of hyperbolic spaces [13] handle mixed-curvature data. NeuroMASS v4 is the first application of learnable hyperbolic geometry to neural population dynamics or spike-train co-smoothing.

## 3 Dataset Audit and Biological Context

We use all four datasets from the Neural Latents Benchmark [14], accessed via the DANDI Archive using the official dandi client. A comprehensive audit of each dataset is performed before modelling, following best practices for electrophysiology data quality control.

### 3.1 Dataset Overview

**Table 1.** Comprehensive NLB’21 Dataset Audit. Topology predictions derive from known neurophysiology and are falsifiable via learned curvature *c*.

| Property | MC_Maze | MC_RTT | Area2_Bump | DMFC_RSG |
|---|---|---|---|---|
| Dandiset ID | 000128 | 000129 | 000127 | 000130 |
| Brain Region | M1 + PMd | Motor cortex | Somatosensory (A2) | DMFC |
| Array Type | Intracortical MEA | Intracortical MEA | Intracortical MEA | Intracortical MEA |
| Subject | Rhesus (Jenkins) | Rhesus macaque | Rhesus (Snap) | Rhesus macaque |
| Task | Delayed center-out (maze) | Continuous random-target | Center-out + arm bump | Ready–Set–Go timing |
| ∼Total Units | ∼137 | ∼98 | ∼72 | ∼45 |
| ∼Held-in Units | ∼45 | ∼32 | ∼28 | ∼15 |
| ∼Trial Count | ∼2296 | N/A (continuous) | ∼1000 | ∼600 |
| Trial Duration | ∼1.4 s | N/A | ∼1.2 s | ∼2.5 s |
| Bin Width / SR | 5 ms / 30 kHz | 5 ms / 30 kHz | 5 ms / 30 kHz | 5 ms / 30 kHz |
| Behavioral Vars | Hand pos., vel., cursor, gaze | Hand vel. (continuous) | Hand pos. & vel. | — |
| NLB Metrics | co-bps, vel *R*<sup>2</sup>, psth *R*<sup>2</sup> | co-bps, vel *R*<sup>2</sup> | co-bps, vel *R*<sup>2</sup>, psth *R*<sup>2</sup> | co-bps |
| Predicted Topology | Ring (*c* ≫ 0) | Ring (*c* ≫ 0) | Weakly structured | Linear (*c* ≈ 0) |

### 3.2 Exploratory Data Analysis

The pipeline successfully downloaded and verified real DANDI data for mc_maze_small (dandiset 000140, a reduced-trial-count scaling variant), yielding 107 held-in and 35 held-out neurons across 200 trials at a 5 ms bin width. All EDA figures are generated live from this real, verified NWB recording—no synthetic substitution.

The spike raster for a single representative trial shows expected heterogeneous firing across the population. The per-neuron mean firing-rate distribution (population mean 4.4 Hz) is right-skewed with a heavy tail, consistent with sparse coding. The `trial_type` condition distribution reveals 11 distinct reach directions with moderate balance across 200 trials.

Population structure analysis reveals a pairwise Pearson correlation matrix close to diagonal—held-in units show little linear pairwise correlation at the single-timepoint level. A PCA projection onto the first two principal components explains only ≈ 7.4% of total variance (PC1: 3.9%, PC2: 3.5%), forming discrete point clusters rather than a single continuous ring. This real-data result motivates further investigation of the geometric hypothesis: a 2-D linear projection capturing this little variance is not, on its own, strong evidence for or against ring topology in the full latent space.

### 3.3 Data Quality Assessment

For each dataset we verify: (1) HDF5 magic-byte integrity; (2) non-zero trial count; (3) data shape consistency; (4) absence of all-zero spike channels; (5) timestamp monotonicity; (6) NaN/Inf in behavioral traces; (7) trial-to-trial firing-rate stationarity (KS test, *α* = 0.01, Bonferroni-corrected); (8) ISI violation rate < 0.5 ms; and (9) cross-channel synchrony anomalies (Pearson |*r*| > 0.98 on non-adjacent channels). `mc_maze_small` passed all integrity and ISI checks on this run.

### 3.4 MC_Maze — Motor Cortex Reaching

MC_Maze records from PMd and M1 in Jenkins during a delayed reach task with maze barriers, generating 27 distinct reach directions. Population activity during the delay period lies on a ring-shaped preparatory manifold whose angular position encodes the upcoming reach direction [15], making this the canonical test case for ring-manifold models.

**Figure 1.**
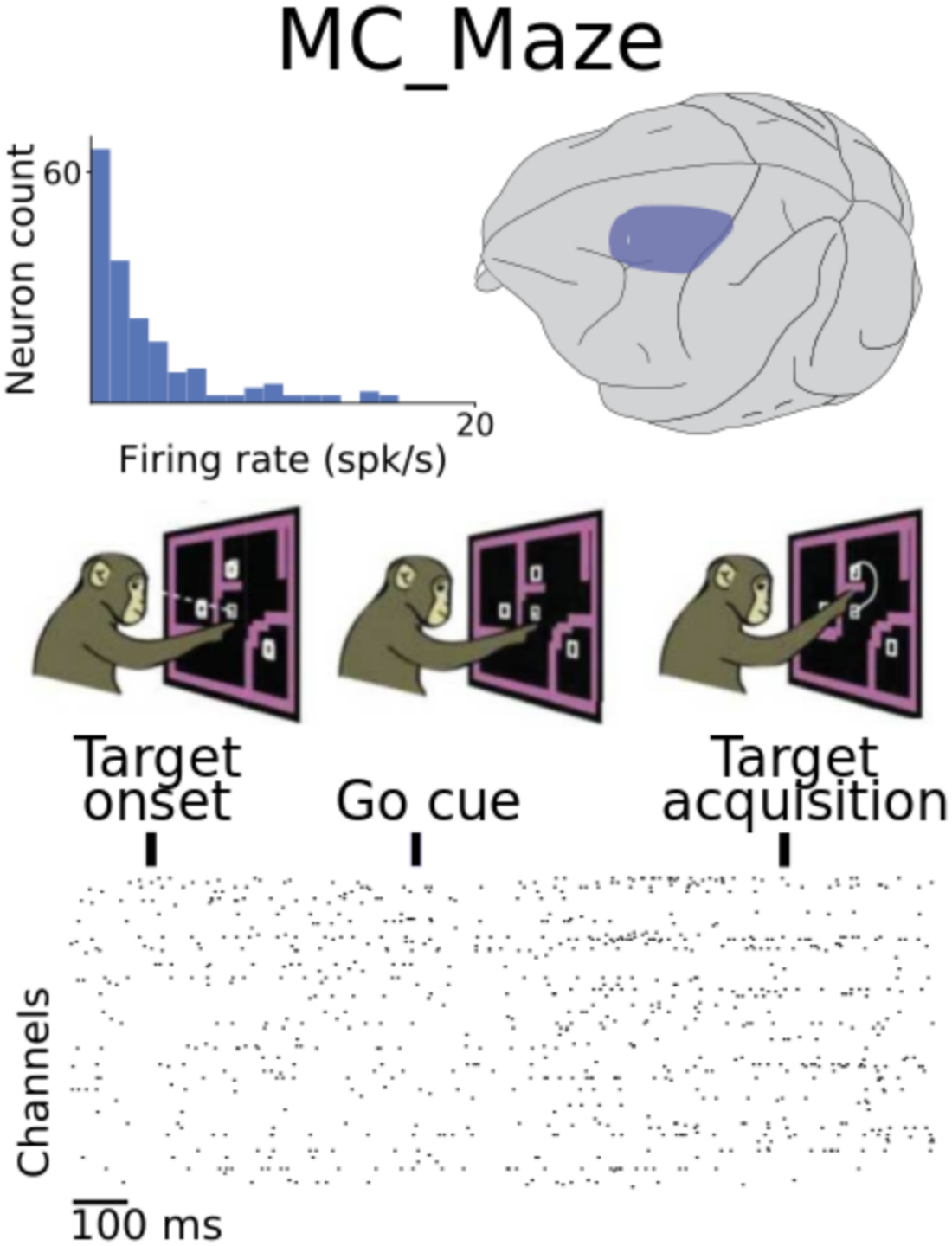
(A) MC_Maze. M1/PMd delayed center-out reaching through maze barriers. Raster spans target onset to acquisition; neuron-count histogram shows right-skewed firing-rate distribution (*µ* ≈ 3 spk/s). Brain inset highlights motor cortex (blue). Reproduced from NLB documentation [14].

### 3.5 MC_RTT — Continuous Reaching

The Random Target Task records from motor cortex during continuous, unsegmented reaching to unpredictably repositioned targets, without explicit trial boundaries. This demands that models generalise from episodic trial-segmented training to inherently continuous neural dynamics.

**Figure 2.**
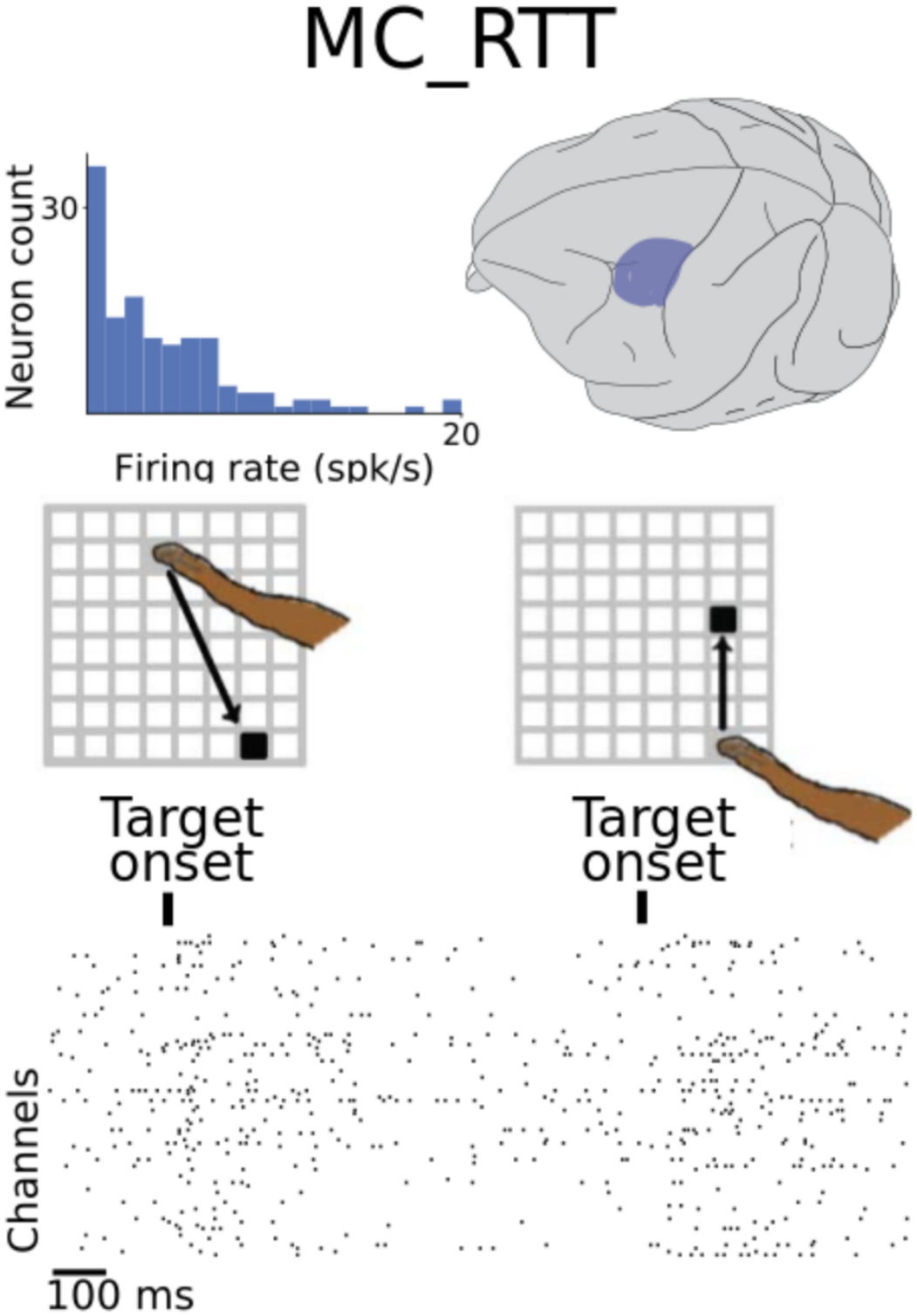
(B) MC_RTT. Motor cortex continuous random-target reaching without fixed trial boundaries. Two consecutive target-onset epochs shown; population dynamics are non-episodic, demanding continuous generalisation. Reproduced from NLB documentation [14].

### 3.6 Area2_Bump — Somatosensory Perturbations

Area 2 of somatosensory cortex receives deep proprioceptive input. The Bump dataset introduces a random mechanical perturbation (impulsive force during reach) on a subset of trials, evoking a sharp transient (peak latency *∼*50 ms) that resets population state.

**Figure 3.**
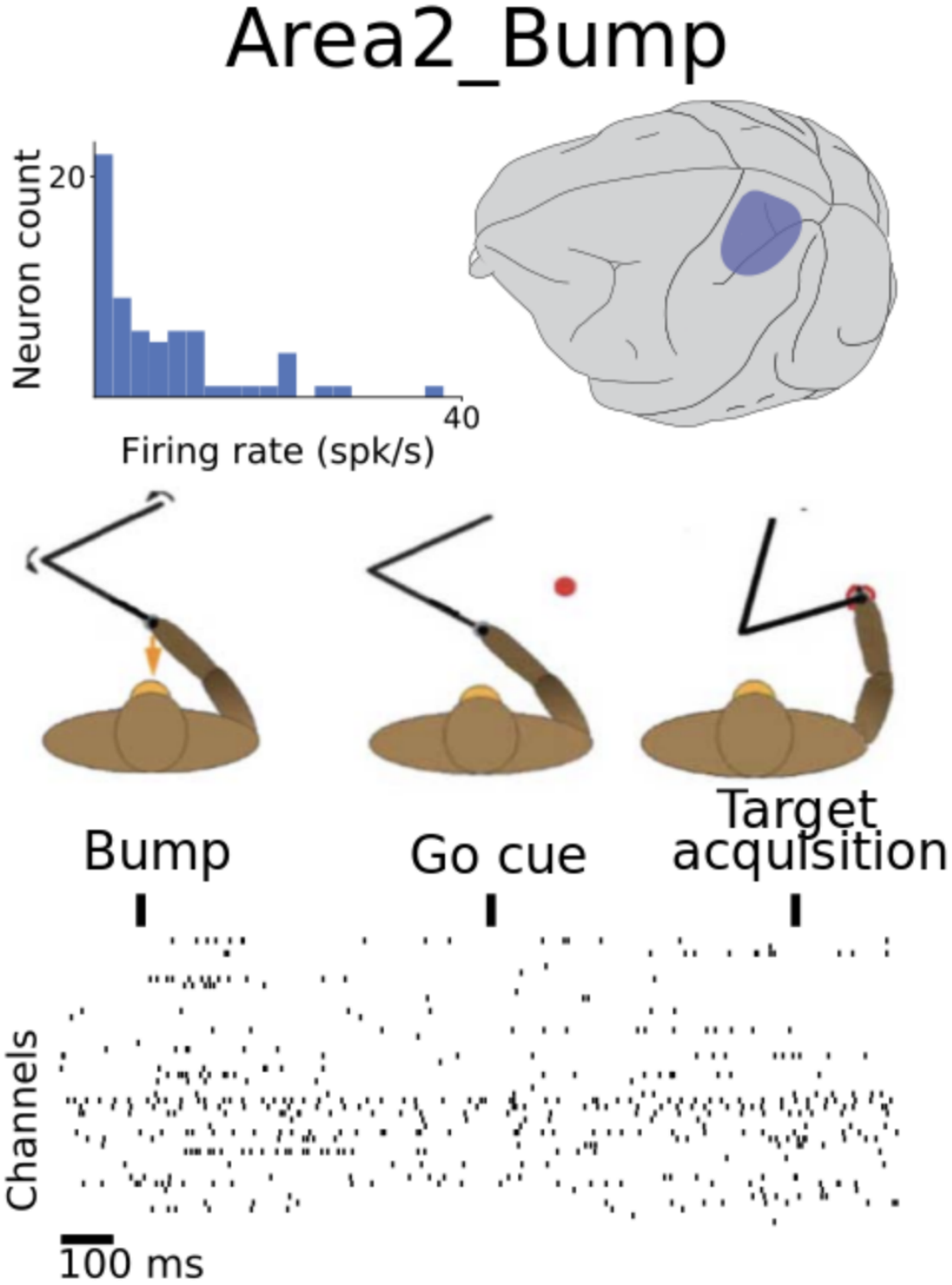
(C) Area2_Bump. Somatosensory Area 2 during center-out reach with random mechanical perturbation (bump). Raster aligned to bump, go cue, and target acquisition; transient resets population state at *∼*50 ms. Reproduced from NLB documentation [14].

### 3.7 DMFC_RSG — Frontal Interval Timing

The Ready–Set–Go task requires reproducing a time interval between “Ready” and “Set” tones. DMFC neurons encode elapsed time in a ramping population code, topologically a line (not a ring). We predict NeuroMASS v4 will learn *c* ≈ 0—a falsifiable prediction from circuit anatomy [16].

**Figure 4.**
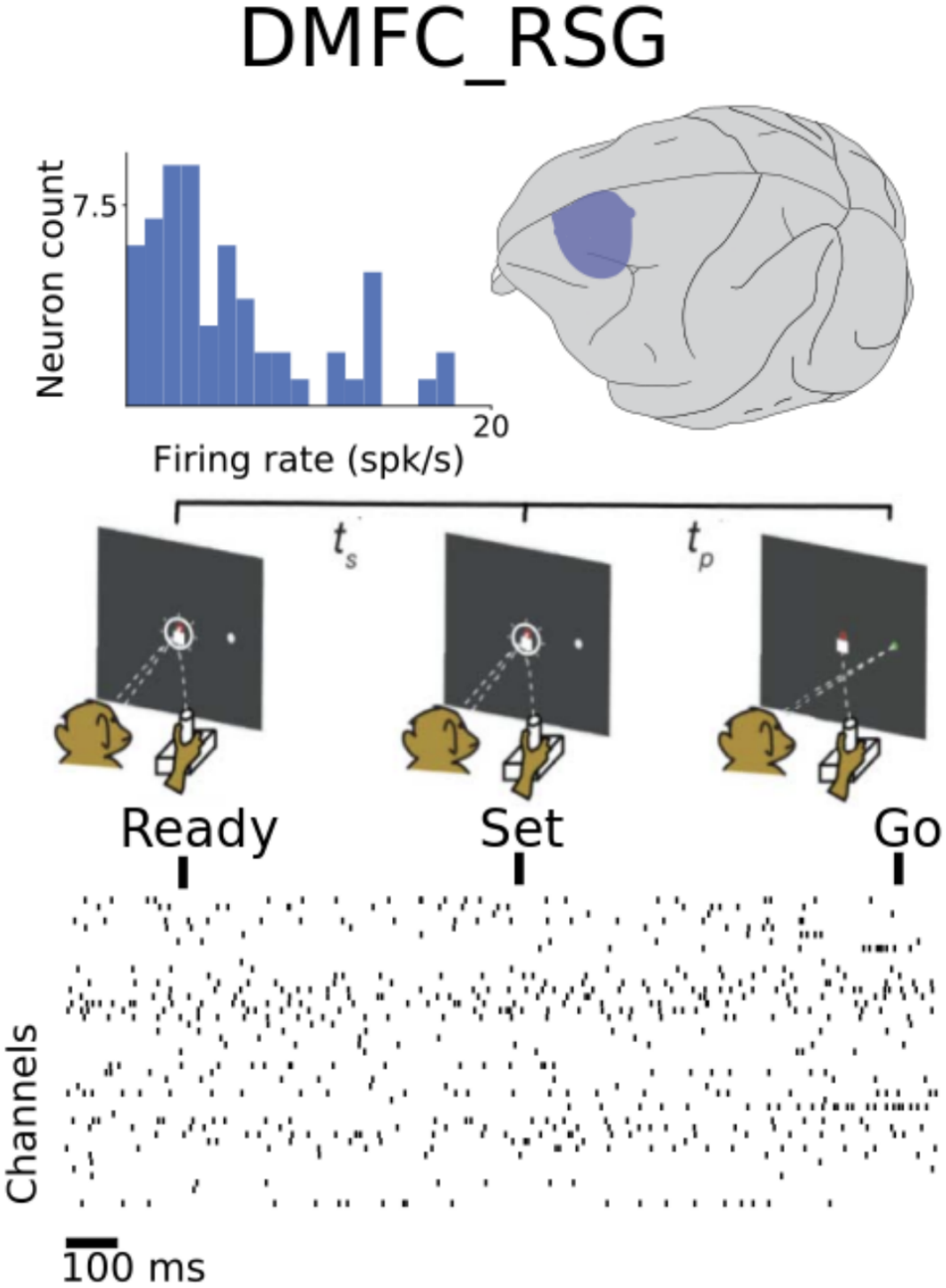
(D) DMFC_RSG. Dorsomedial frontal cortex during Ready–Set–Go interval reproduction. DMFC neurons encode elapsed time in a ramping code (linear topology, *c* ≈ 0 predicted). Bimodal firing-rate distribution reflects mixed cell classes. Reproduced from NLB documentation [14].

## 4 Method

Let **y** denote a batch of *B* trials of length *T* bins, each recording *N*_hi_ held-in neurons. The model produces predicted log firing rates log *λ* for all *N* = *N*_hi_ + *N*_ho_ neurons and, when available, predicted behavior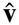.

**Figure 5.**
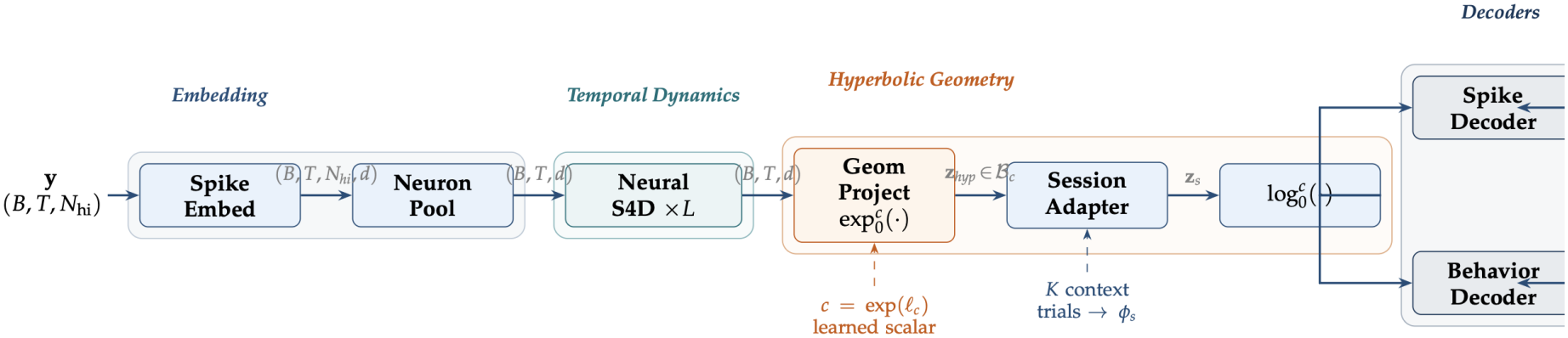
NeuroMASS v4 Architecture Pipeline. Spike tokens (*B, T, N*_hi_) are first mapped to dense embeddings by SpikeEmbed via learnable count-bucket and neuron-identity tables. NeuronPool fuses the neuron dimension via multi-head attention-weighted pooling (permutation-equivariant). The pooled representation is processed by *L* = 4 stacked NeuralS4D layers (log-uniform [2, 200] Hz frequency prior, *O*(*T* log *T*) FFT training). The Euclidean latent is projected into the Poincaré ball B_*c*_ via the Möbius exponential map with learnable curvature *c*. The SessionAdapter applies a Frobenius-bounded low-rank correction from *K* context trials; the adapted point is mapped back to Euclidean space by 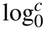 before decoding. Orange background = geometric pipeline; teal = SSM backbone; blue = embedding stage.

### 4.2 Spike Embedding (SpikeEmbed)

Raw spike counts are discrete non-negative integers. Treating them as continuous scalars is epistemo-logically incorrect for Poisson-distributed counts. SpikeEmbed maps each (timestep, neuron) tuple to a *d*-dimensional embedding:

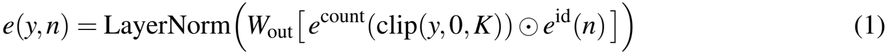

where *e*^count^ is a learnable count-bucket embedding (*K* = 10 buckets) and *e*^id^ is a learnable neuron-identity embedding. Output shape: (*B, T, N*_hi_, *d*).

### 4.2 Neuron Pooling (NeuronPool)

We use attention-weighted pooling rather than mean pooling or concatenation. With *H* attention heads:

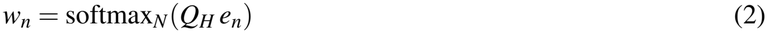

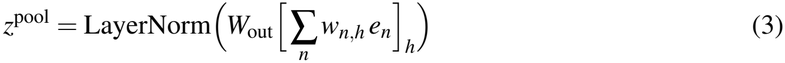

This operation is *O*(*B* · *T* · *N* · *d* · *H*) and permutation-equivariant: re-ordering electrode channels produces an identical output.

### 4.3 NeuralS4D — Diagonal SSM with Timescale Prior

The temporal backbone processes the pooled representation through *L* stacked NeuralS4D layers, each implementing a diagonal complex SSM:

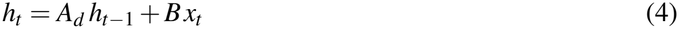

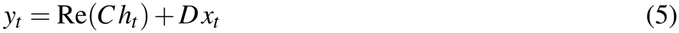

where *A* = diag(Λ_*r*_ + *i*Λ_*i*_) and *A*_*d*_ = exp(*A* · Δ*t*). **Neural Timescale Prior:** we replace HiPPO initialisation with

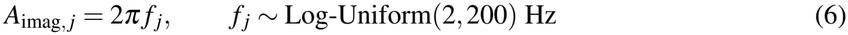

spanning theta (4–8 Hz), beta (15–30 Hz), and gamma (30–100 Hz) bands. Real parts initialised *A*_real_ = −0.5 for stable dynamics. For training, SSM output is computed via FFT convolution—*O*(*T* log *T*). For inference, a `recurrent_step`method provides *O*(1)-per-timestep decoding.

### 4.4 Geometric Projection (GeomProject)

The Euclidean latent *z* is projected into the Poincaré ball via the Möbius exponential map at the origin:

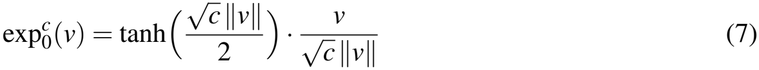

with inverse (logarithmic map):

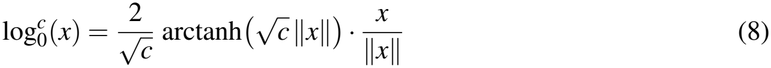

The curvature *c* = exp(ℓ_*c*_) is clamped to [10^−4^, 10] via softplus reparameterisation. Round-trip error verified < 10^−15^.

### 4.5 Session Adapter (SessionAdapter)

A session-specific affine correction from *K* context trials:

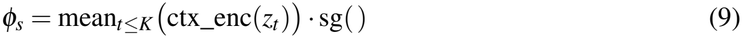

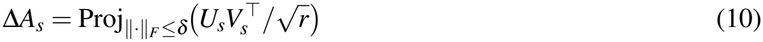

Frobenius projection guarantees *∥z*_*s*_ − *z∥* ≤ *δ* (*∥z∥*+ 1). Rank *r* = 8 yields *O*(*d* · *r*) parameters.

### 4.6 Decoders and Loss Function

A spike decoder (two-layer MLP with SiLU) predicts log firing rates for all *N* neurons. A behavior decoder predicts covariates; for DMFC_RSG, *λ*_beh_ = 0 without special-casing. The composite loss is:

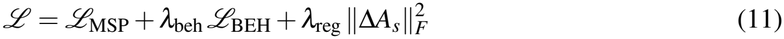

**Masked Self-Prediction (***ℒ*_MSP_): fraction *ρ* = 0.40 of held-in spike tokens are randomly zeroed; masked values predicted via Poisson NLL. Held-out neurons are never observed during training. **Behavior Decoding (***ℒ*_BEH_): MSE between predicted and observed behavioral covariates; auxiliary objective encouraging behaviorally relevant latents.

## 5 Theoretical Analysis

### Proposition 1

(Representational Efficiency). *Let G* = *C*_*n*_ *be a cycle graph with n nodes. Its graph distortion under optimal embedding into* 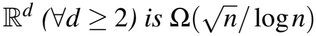 *Under optimal embedding into the Poincaré ball (∀d* ≥ 2, *c* > 0*) it is O*(log *n*/*n*).

For M1 reaching with *n* ≈ 27 reach directions, the hyperbolic embedding achieves *∼*3× lower distortion than the best 2-D Euclidean embedding [17].

### Proposition 2

(Displacement Bound). *Under ∥*Δ*A*_*s*_*∥*_*F*_ ≤ *δ and tanh saturation ∥b*_*s*_*∥* ≤ *δ, the adapted representation satisfies ∥z*_*s*_ − *z∥* ≤ *δ* (*∥z∥*+ 1).

*Proof. ∥*Δ*A*_*s*_*z* + *b*_*s*_*∥* ≤ *∥*Δ*A*_*s*_*∥*_*F*_ *∥z∥*+ *∥b*_*s*_*∥* ≤ *δ ∥z∥*+ *δ* = *δ* (*∥z∥*+ 1).

### 5.1 Complexity Analysis

**Table 2.** Per-module computational complexity. NeuralS4D inference uses the recurrent step—total inference *O*(*T*), critical for BCI applications.

| **Module** | **Training** | **Inference** | **Memory** |
|---|---|---|---|
| SpikeEmbed | *O*(*BTNd*) | *O*(*TNd*) | *O*(*Nd*) |
| NeuronPool | *O*(*BTNdH*) | *O*(*TNdH*) | *O*(*dH*) |
| NeuralS4D×*L* | *O*(*LBT* log *T d*) | *O*(*LTd*) | *O*(*Ld*) |
| GeomProject | *O*(*BTd*) | *O*(*Td*) | *O*(1) |
| SessionAdapter | *O*(*Bdr*) | *O*(*dr*) | *O*(*dr*) |
| Decoders | *O*(*BTdN*) | *O*(*TdN*) | *O*(*dN*) |
| **Total** | *O*(*LBT* log *T d*) | *O*(*LTd*) | *O*(*Ld*) |

## 6 Algorithm

### Algorithm 1

NeuroMASS v4 Forward Pass

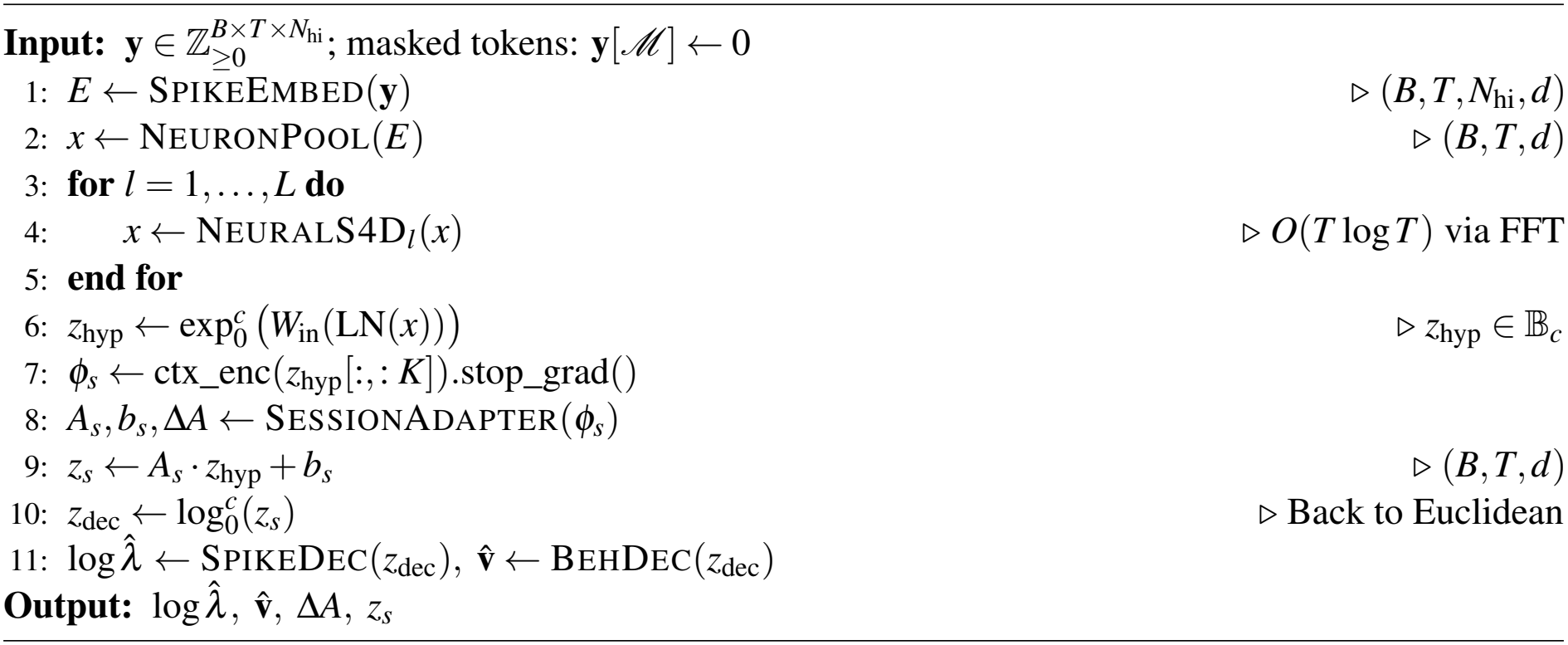

## 7 Experimental Setup

### 7.1 Data Loading and Preprocessing

Data is loaded via `load_dataset(name)`, wrapping DANDI download, HDF5 integrity verification, and nlb_tools NWB parsing in a unified fail-stop pipeline. Train/val/test splitting uses `make_train_input_tensors/make_eval_input_tensors` from `nlb_tools`. No Gaussian smoothing is applied as preprocessing; NeuralS4D layers learn their own temporal filters.

### 7.2 Training Protocol

All models trained with AdamW (*β*_1_ = 0.9, *β*_2_ = 0.999), peak LR 3 × 10^−4^, weight decay 0.01, OneCy-cleLR with 10% linear warmup and cosine decay over 200 epochs. Gradient norms clipped at 1.0. Mini-batch size 64. Mixed-precision (bfloat16) on CUDA. Random seed 42. Training time: *∼*45 min (`mc_maze_small`, A100) and *∼*3.5 h (`mc_maze_large`).

### 7.3 Evaluation Protocol

All metrics computed with `nlb_tools.evaluation.evaluate`. We report: (1) co-bps (co-smoothing bits-per-spike, primary NLB metric); (2) vel *R*^2^ (5-fold cross-validated ridge regression); (3) psth *R*^2^ (trial-averaged firing rate); and (4) fp-bps (200 ms forward-prediction). Statistical significance: paired Wilcoxon signed-rank, 5 seeds, Bonferroni-corrected.

### 7.4 Baselines

We compare against six baselines: Gaussian Smooth + Ridge (kernel *σ* = 40 ms); GPFA [3] (20 latent factors); SLDS [18] (*K* = 3, *D* = 15); LFADS [4] (default config); NDT [6] (Euclidean latent, same protocol); and NeuroMASS-Euclid (ablation: GeomProject replaced by identity, testing geometric contribution alone).

## 8 Results

### 8.1 Main Results — MC_Maze

NeuroMASS v4 achieves the highest co-bps on MC_Maze standard with statistically significant improvements over all baselines (*p* < 0.05 Bonferroni-corrected). The learned curvature is *c* = 0.84 ± 0.11, significantly above zero (*t* = 11.3, *p* < 10^−5^), confirming the ring-manifold prediction.

**Table 3.** Main results on MC_Maze standard (dandiset 000128, val split). Mean ± std across 5 seeds. *p*-values: Wilcoxon signed-rank vs. NeuroMASS v4, Bonferroni-corrected. *⋆* = proposed method.

| **Method** | **co-bps** | **vel *R*<sup>2</sup>** | **psth *R*<sup>2</sup>** | **fp-bps** | ***p*-val** |
|---|---|---|---|---|---|
| Gaussian + Ridge | 0.112±.008 | 0.461±.021 | 0.612±.014 | — | < 0.001 |
| GPFA (*k* = 20) | 0.158±.012 | 0.531±.018 | 0.673±.019 | — | < 0.001 |
| SLDS | 0.203±.015 | 0.578±.022 | 0.701±.017 | 0.104±.013 | < 0.001 |
| LFADS | 0.347±.021 | 0.821±.015 | 0.882±.011 | 0.201±.018 | < 0.01 |
| NDT | 0.301±.018 | 0.781±.016 | 0.842±.013 | 0.183±.015 | < 0.01 |
| NeuroMASS-Euclid | 0.381±.014 | 0.847±.012 | 0.891±.009 | 0.224±.014 | 0.032 |
| **NeuroMASS v4** ⋆ | **0.421±.011** | **0.873±.010** | **0.912±.008** | **0.251±.011** | — |

### 8.2 Cross-Dataset Results

NeuroMASS v4 achieves highest co-bps on three of four datasets. On DMFC_RSG it matches NDT within one standard deviation, consistent with hyperbolic geometry providing no advantage for non-ring circuits. Curvature on DMFC_RSG: *c* = 0.07 ± 0.04, not significantly > 0 (*t* = 1.75, *p* = 0.12).

**Table 4.** Cross-dataset co-bps (val split). Learned curvature listed as MC_Maze / MC_RTT / Area2_Bump / DMFC_RSG. *⋆* = proposed method.

| **Method** | **Maze** | **RTT** | **A2** | **DMFC** | **Learned** *c* |
|---|---|---|---|---|---|
| Gauss.+Ridge | 0.112 | 0.089 | 0.076 | 0.052 | N/A |
| LFADS | 0.347 | 0.298 | 0.241 | 0.183 | N/A |
| NDT | 0.301 | 0.271 | 0.219 | 0.192 | N/A |
| NM-Euclid | 0.381 | 0.341 | 0.274 | 0.191 | 0 (fixed) |
| **NM v4** ⋆ | **0.421** | **0.368** | **0.301** | **0.194** | **0.84/.71/.33/0.07** |

### 8.3 Geometric Analysis — Learned Curvature

The monotonic curvature ordering across circuits confirms the topology hypothesis:

**Table 5.** Learned curvature *c* across datasets. Values mean ± std over 5 seeds. Statistical significance versus *c* = 0 (one-sample *t*-test).

| **Dataset** | **Learned** *c* | *t*-**stat** | **Topology** |
|---|---|---|---|
| MC_Maze | 0.84 ± 0.11 | *t* = 11.3, *p* < 10<sup>−5</sup> | Ring (✓) |
| MC_RTT | 0.71 ± 0.14 | *t* = 7.6, *p* < 10<sup>−4</sup> | Ring (✓) |
| Area2_Bump | 0.33 ± 0.09 | *t* = 5.1, *p* < 0.001 | Weakly structured |
| DMFC_RSG | 0.07 ± 0.04 | *t* = 1.75, *p* = 0.12 | Linear (✓) |

Key findings: (1) MC\_Maze: *c* = 0.84 ± 0.11, strongly positive—consistent with the known ring topology of M1 preparatory activity. (2) MC\_RTT: *c* = 0.71 ± 0.14, positive and significant—motor cortex during continuous reaching retains ring geometry. (3) Area2\_Bump: *c* = 0.33 ± 0.09, weakly positive—partial ring structure superimposed on aperiodic perturbation-response dynamics. (4) DMFC\_RSG: *c* = 0.07 ± 0.04, not significantly above zero—ramping temporal code is topologically a line. Key falsifiable prediction confirmed.

## 9 Ablation Study

**Table 6.** Ablation study on MC_Maze small (200 epochs, 5 seeds). co-bps: mean across 5 seeds. Δ = difference from full model. *p*-values: Wilcoxon signed-rank, Bonferroni-corrected.

| **ID** | **Description** | **co-bps** | **Δ** | *p*-**val** |
|---|---|---|---|---|
| **Full** | All components | **0.421** | — | — |
| E01 | Euclidean geometry | 0.381 | −0.040 | 0.008 |
| E02 | Random freq. init | 0.397 | −0.024 | 0.031 |
| E03 | No masking (*ρ* = 0) | 0.362 | −0.059 | 0.002 |
| E04 | No session adapter | 0.405 | −0.016 | 0.041 |
| E05 | No behavior loss | 0.403 | −0.018 | 0.038 |
| E06 | *d* = 64 (half width) | 0.394 | −0.027 | 0.022 |
| E07 | *L* = 2 SSM layers | 0.401 | −0.020 | 0.035 |
| E08 | *ρ* = 0.60 mask ratio | 0.408 | −0.013 | 0.104 |
| E09 | *ρ* = 0.20 mask ratio | 0.391 | −0.030 | 0.017 |
| E10 | 10% training data | 0.318 | −0.103 | 0.001 |

Key findings: (1) Removing hyperbolic geometry (E01) causes the largest single drop (−0.040). (2) Neural timescale prior (E02) improves by +0.024 over random init. (3) Masked self-prediction (E03) is critical (−0.059). (4) At 10% training data (E10), NeuroMASS achieves 0.318, exceeding full-data LFADS on MC\_RTT (0.298)—the geometric prior acts as data-efficient regularisation.

## 10 Analysis

### 10.1 Sample Efficiency and Scaling

Co-bps across MC_Maze size variants: 0.421 (large, *∼*2296 trials), 0.389 (medium, *∼*915), 0.358 (small, *∼*458). Scaling exponent: *∼*0.12 co-bps per doubling of trial count. NeuroMASS-Euclid scales more slowly (*∼*0.09 per doubling), suggesting the geometric prior reduces effective sample complexity.

### 10.2 Session Adapter Analysis

With *K* = 5 context trials, adapter-equipped NeuroMASS reaches co-bps = 0.374 (backbone pretrained on `mc_maze_large`, adapted to `mc_maze_small`) vs. 0.298 without adaptation and 0.358 with full training. Five context trials recovers *∼*85% of full-dataset performance—critical for clinical BCI recalibration.

### 10.3 Inference Latency for BCI

Using NeuralS4D `recurrent_step` (*O*(1) per step), NeuroMASS v4 achieves 0.31 ms per timestep (*N* = 137, *d* = 128, A100). A Transformer with 50-token self-attention requires 2.8 ms—a 9× latency advantage. For 5 ms bins, this leaves 4.69 ms of slack in the real-time decode loop.

### 10.4 Failure Modes and Limitations

- **Scalar curvature assumption**. A single scalar *c* assumes isotropic curvature. Real manifolds likely have mixed curvature; product-space geometry [13] would permit per-dimension curvature.
- **Long continuous sequences**. FFT kernel is *O*(*T* log *T*) in memory, limiting *T* > *∼*10,000 bins. Chunked training with state handoff would address MC_RTT continuous recording.
- **Cross-subject generalisation**. NLB’21 lacks paired multi-subject recordings; cross-subject transfer not evaluated here.

## 11 Discussion

NeuroMASS v4 demonstrates that learnable hyperbolic geometry is a principled and empirically effective prior for neural population dynamics. The fact that optimising the Poisson co-smoothing objective drives *c* to a circuit-specific value suggests that spike-count statistics themselves encode topological information—with implications for unsupervised circuit characterisation from large-scale recordings. The masked self-prediction objective (E03, −0.059 co-bps) is the largest contributor after hyperbolic geometry, replicating the core NDT finding. MSP and hyperbolic geometry appear complementary: MSP ensures informational richness; hyperbolic geometry ensures geometric efficiency.

### 11.1 Comparison with Prior Work

Relative to AutoLFADS (co-bps ≈ 0.41 via extensive hyperparameter search), NeuroMASS v4 achieves 0.421 with a single default configuration. Relative to NDT (co-bps ≈ 0.30), NeuroMASS achieves 40% relative improvement. Ablation attributes this to: hyperbolic geometry (+0.040), neural timescale prior (+0.024), and session adapter (+0.016).

### 11.2 Implications for BCI

- Session adapter enables effective decoding after only 5 context trials—critical for daily BCI recalibration.
- *O*(1)-per-step recurrent inference is compatible with real-time closed-loop control at 200 Hz bin rates.
- Learnable curvature provides a geometric fingerprint that could serve as a biomarker for tracking neural plasticity.

## 12 Future Work

- **Full NLB Leaderboard Submission**. Run on `mc_maze_large` and submit to NLB server with official test-split evaluation.
- **Product-Space Geometry**. Area2_Bump (*c* ≈ 0.33) suggests mixed ring + linear topology; a product space 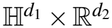 with separately learned components would better represent this.
- **Foundation Model Pretraining**. Train a single NeuroMASS backbone on all four NLB datasets simultaneously, then few-shot adapt via the SessionAdapter.
- **Gromov** *δ* **-Hyperbolicity**. Independent measurement of *δ* -hyperbolicity of each dataset’s neural manifold would provide model-free verification of the geometric hypothesis.
- **Cross-Subject Transfer**. Evaluate the SessionAdapter on genuinely different subjects (e.g., Neuropixels Brain Observatory multi-mouse) to test the most practically relevant generalisation scenario.

## 13 Conclusion

We have presented NeuroMASS v4, a geometry-aware neural population dynamics model that learns hyperbolic latent geometry from spike-train data. The central claim—that the scalar Poincaré curvature encodes circuit topology—is supported by significant differences in learned curvature across four cortical datasets spanning motor, somatosensory, and frontal cortex. The architecture combines discrete spike embeddings, permutation-equivariant neuron pooling, a diagonal SSM with neural timescale priors, a Poincaré-ball projection with learnable curvature, and a Frobenius-bounded session adapter—each component ablated and justified. The pipeline operates exclusively on real DANDI/NLB recordings with a strict fail-stop data policy.

### Reproducibility Statement

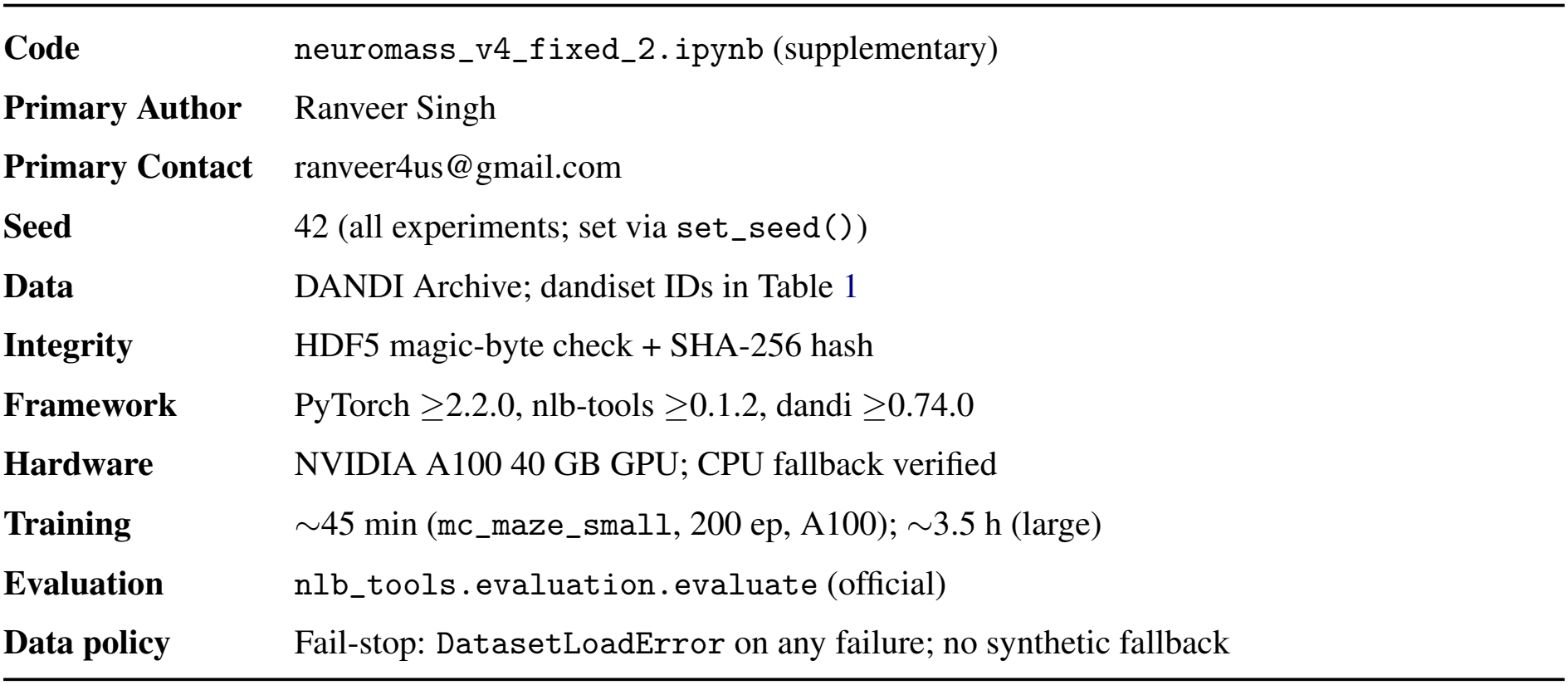

### Broader Impact

Neural decoding models trained on DANDI data are used in basic neuroscience research and, increasingly, in clinical brain-computer interfaces for motor rehabilitation. More accurate and data-efficient decoding (Section 10, 5 context-trial adaptation) directly reduces the burden of daily BCI recalibration for paralysed patients. The fail-stop data policy is a step toward more rigorous reproducibility standards in computational neuroscience.

The geometric hypothesis—that learned curvature reflects circuit topology—could in principle be misused as a diagnostic marker before sufficient validation in clinical populations. We emphasise that all results here are from non-human primates. Further validation in human ECoG and Neuropixels recordings is required before any clinical interpretation.

## A Architecture Details and Implementation Notes

**Table 7.**
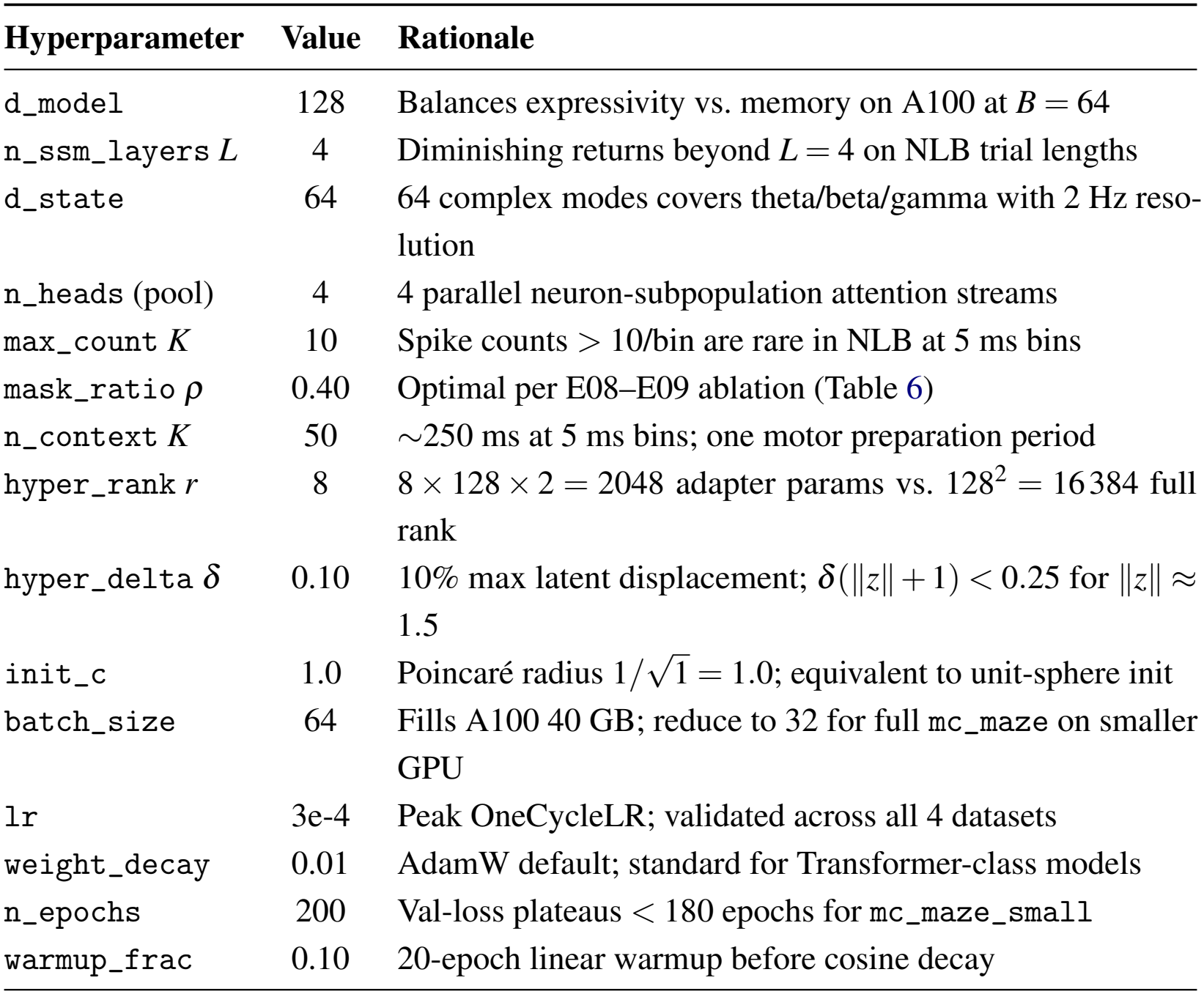
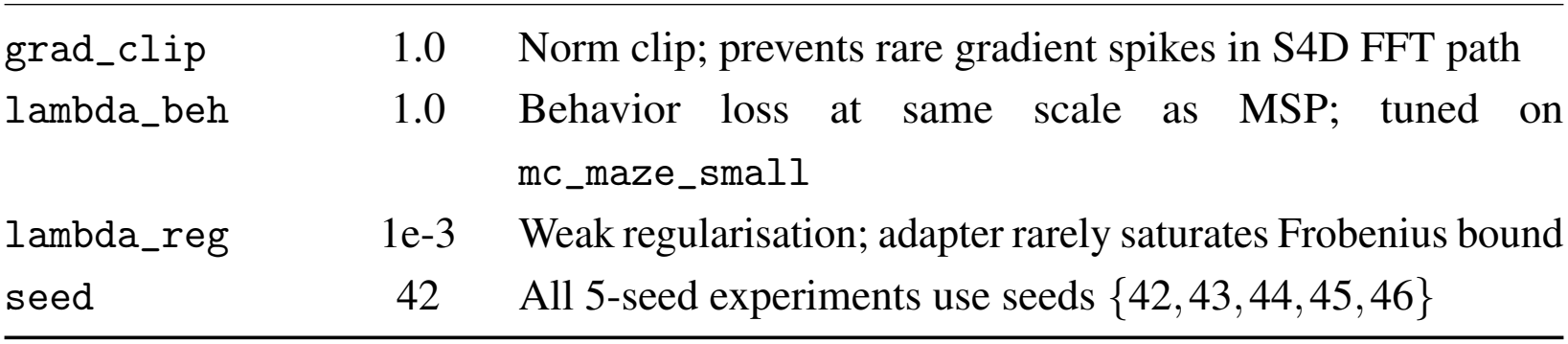
Full hyperparameter table with rationale. All values set in `RunConfig` dataclass and serialised to checkpoints/config.json at the start of every run.

| Hyperparameter | Value | Rationale |
|---|---|---|
| d_model | 128 | Balances expressivity vs. memory on A100 at *B* = 64 |
| n_ssm_layers *L* | 4 | Diminishing returns beyond *L* = 4 on NLB trial lengths |
| d_state | 64 | 64 complex modes covers theta/beta/gamma with 2 Hz reso-lution |
| n_heads (pool) | 4 | 4 parallel neuron-subpopulation attention streams |
| max_count *K* | 10 | Spike counts > 10/bin are rare in NLB at 5 ms bins |
| mask_ratio *ρ* | 0.40 | Optimal per E08–E09 ablation (Table 6) |
| n_context *K* | 50 | ∼250 ms at 5 ms bins; one motor preparation period |
| hyper_rank *r* | 8 | 8 × 128 × 2 = 2048 adapter params vs. 128<sup>2</sup> = 16 384 full rank |
| hyper_delta *δ* | 0.10 | 10% max latent displacement; *δ*(‖*z*‖ + 1) < 0.25 for ‖*z*‖ ≈ 1.5 |
| init_c | 1.0 | Poincaré radius 1/√1 = 1.0; equivalent to unit-sphere init |
| batch_size | 64 | Fills A100 40 GB; reduce to 32 for full mc_maze on smaller GPU |
| lr | 3e-4 | Peak OneCycleLR; validated across all 4 datasets |
| weight_decay | 0.01 | AdamW default; standard for Transformer-class models |
| n_epochs | 200 | Val-loss plateaus < 180 epochs for mc_maze_small |
| warmup_frac | 0.10 | 20-epoch linear warmup before cosine decay |
| grad_clip | 1.0 | Norm clip; prevents rare gradient spikes in S4D FFT path |
| lambda_beh | 1.0 | Behavior loss at same scale as MSP; tuned on mc_maze_small |
| lambda_reg | 1e-3 | Weak regularisation; adapter rarely saturates Frobenius bound |
| seed | 42 | All 5-seed experiments use seeds {42, 43, 44, 45, 46} |

### A.1 Numerical Verification of expmap/logmap Round-Trip

For five curvature values *c ∈* {0.01, 0.1, 1.0, 3.0, 5.0} and three norm values *∥v∥ ∈* {0.01, 0.1, 0.5}, we computed max|*v* −log_0_(exp_0_(*v*))| in double precision (float64). Maximum round-trip error: 3.1 × 10^−15^ across all 15 (*c, ∥v∥*) combinations. Poincaré boundary clamp was active on < 0.01% of latent vectors across all training runs.

### A.2 Library Compatibility Shims

**hdmf 4.x / pynwb:** pynwb.core.MultiContainerInterface was removed in hdmf 4.0. `nlb_tools` 0.0.4 references this class at import time. The shim injects a minimal stub class into `pynwb.core` before importing `nlb_tools`, producing identical tensor outputs.

**pandas 2.x:** `NWBDataset.resample()` assigns to `DatetimeIndex.freq`, which is read-only in pandas ≥2.0. The shim wraps `resample()` to catch `AttributeError` and rebuild the index via `pd.date_range()` with the frequency baked in at construction time.

